# The metalloproteinase inhibitor Marimastat improves skeletal muscle regeneration when administered intravenously after myonecrosis induced by the venom of *Bothrops asper*

**DOI:** 10.64898/2026.03.25.714270

**Authors:** Andrés Zamora, Alexandra Rucavado, Teresa Escalante, José María Gutiérrez, Erika Camacho

**Affiliations:** Instituto Clodomiro Picado, Facultad de Microbiología; Departamento de Fisiología, Escuela de Medicina, Universidad de Costa Rica, San José 11501, Costa Rica

**Keywords:** *Bothrops asper* venom, myonecrosis, muscle regeneration, fibrosis, Varespladib, Marimastat

## Abstract

Skeletal muscle regeneration is often impaired after acute muscle damage induced by viperid snake venoms, such as that of *Bothrops asper*, a medically-relevant species in Latin America. It has been shown that traces of venom that remain in the damaged muscle affect myogenic cells in culture, raising the possibility of inhibition of these toxins during the regenerative process as a way to improve regeneration. Using a mouse model of myonecrosis and regeneration, we evaluated the effects of Varespladib (a phospholipase A_2_ inhibitor) or Marimastat (a metalloproteinase inhibitor) on muscle regeneration when administered intravenously 24 h after the onset of myonecrosis, i.e., after muscle damage has occurred. The regenerative process was evaluated 14 and 28 days after venom injection. Results show that Marimastat, or a combination of both inhibitors, improved the extent of skeletal muscle regeneration and reduced the extent of tissue fibrosis when compared to tissue from mice receiving venom and no inhibitors, as judged by qualitative and quantitative histological assessment. Results underscore the deleterious role of traces of venom components in the damaged muscle during muscle regeneration and suggest that the administration of metalloproteinase inhibitors, or a combination of metalloproteinase and phospholipase A_2_ inhibitors, even when muscle damage has developed, may be a therapeutic alternative for improving the extent of muscle regeneration.

## 1. Introduction

Envenomings by viperid snake species often result in pronounced damage to skeletal muscle, i.e., myonecrosis [1–3]. Despite the fact that skeletal muscle has an intrinsic capacity to regenerate after several types of injury [4], in the case of viperid venoms, such as that of *Bothrops asper*, the regenerative response is often impaired [5–7], with the consequent loss of muscle mass and its replacement by fibrotic and adipose tissues, resulting in long-term sequelae which drastically affect the quality of life of snakebite victims [1,8]. The reasons behind such poor regenerative outcome are not completely understood, but are likely to be multifactorial, involving damage to the microvasculature and intramuscular nerves, deficient removal of necrotic debris, and degradation of the basement membrane of muscle fibers, among other mechanisms [5–7, 9].

When investigating the effect of homogenates of skeletal muscle obtained from mice after myonecrosis induced by *B. asper* venom or isolated muscle-damaging toxins, it was observed that these homogenates inhibited myoblast replication and fusion into myotubes in cell culture even when the homogenates were from muscles obtained five days after venom injection [10], an effect also observed with the venoms of other *Bothrops* species [11]. Since such effects were abrogated when the homogenates were incubated with antivenom, it was concluded that traces of venom in the regenerating tissue were responsible for affecting muscle myogenic cells. Moreover, the venoms of the viperids *Daboia russelii* and *Bitis arietans* are also known to affect myoblast viability, to inhibit their fusion into myotubes, and to induce atrophy of myotubes in culture [12,13]. Thus, traces of venom remaining in muscle tissue after myonecrosis may affect regenerating muscle cells and contribute to the impairment of the regenerative process in envenomings by a variety of viperid species, raising the possibility of using inhibitors of muscle damaging toxins along the regenerative process after acute myonecrosis as a therapeutic tool to enhance regeneration.

Since myotoxic phospholipases A_2_ (PLA_2_s) and hemorrhagic metalloproteinases (SVMPs) play key roles in acute muscle damage induced by *B. asper* venom [14], and homogenates from muscle tissue injected with these toxins impaired myoblast fusion to form myotubes [10], we hypothesized that the administration of inhibitors of these enzymes, i.e., Varespladib and Marimastat, after the development of local myonecrosis, may result in an improved skeletal muscle regeneration. In this report we show that muscle regeneration improves when Marimastat or a combination of Marimastat and Varespladib are administered 24 h after the onset of *B. asper* venom-induced myonecrosis.

## 2. Results

### 2.1. Muscle necrosis induced by B. asper venom

To assess the extent of skeletal muscle damage induced by the injection of 50 µg *B. asper* venom in the gastrocnemius muscle, tissue sections were analyzed 24 h after venom injection, since previous studies demonstrated that tissue damage induced by *B. asper* venom occurred within the first hours after injection [15–17]. Thus, assessing tissue necrosis at 24 h allowed the quantification of the net extent of muscle damage. Widespread necrosis of muscle fibers, extravasated erythrocytes and abundant inflammatory cells were observed (Supplementary figure S1), as characteristic pathological alterations induced by this venom [17]. The muscle necrotic area, characterized by hypercontracted muscle fibers, corresponded to 48.2 ± 13.7 % of the total area of muscle. This observation agrees with the value of 50% necrosis found in a previous study using the same experimental model with this venom [16]. With this basis we then assessed the effect of inhibitors on skeletal muscle regeneration when administered 24 h after venom.

### 2.2. Effect of Varespladib and Marimastat on skeletal muscle regeneration after necrosis induced by B. asper venom

#### 2.2.1. Observations in samples collected 14 days post-envenoming

Tissue sections collected 14 days after injection of PBS showed a normal histological pattern of muscle tissue, without fibrosis and devoid of muscle fibers with centrally located nuclei. On the other hand, at this time interval tissue of mice that received *B. asper* venom and then PBS instead of inhibitors showed scarce and dispersed areas of muscle regeneration, characterized by small muscle fibers with centrally-located muscle, and areas where fibrosis and granulation tissue predominated (Fig 1). In contrast, sections from muscle of mice that were treated with either Varespladib or Marimastat 24 h after venom injection showed a histological pattern characterized by a higher number of regenerating muscle fibers with centrally-located nuclei and less areas of poor regeneration. This trend was more pronounced in samples from envenomed mice treated with the combination of Varespladib and Batimastat (Fig 1). Tissue sections stained with picrosirius red evidenced a high extent of fibrosis in samples from envenomed mice not receiving the inhibitors (Fig 1), whereas less abundant picrosirius red-stained areas were observed in samples of envenomed mice receiving the inhibitors (Fig 1).

**Figure 1.**
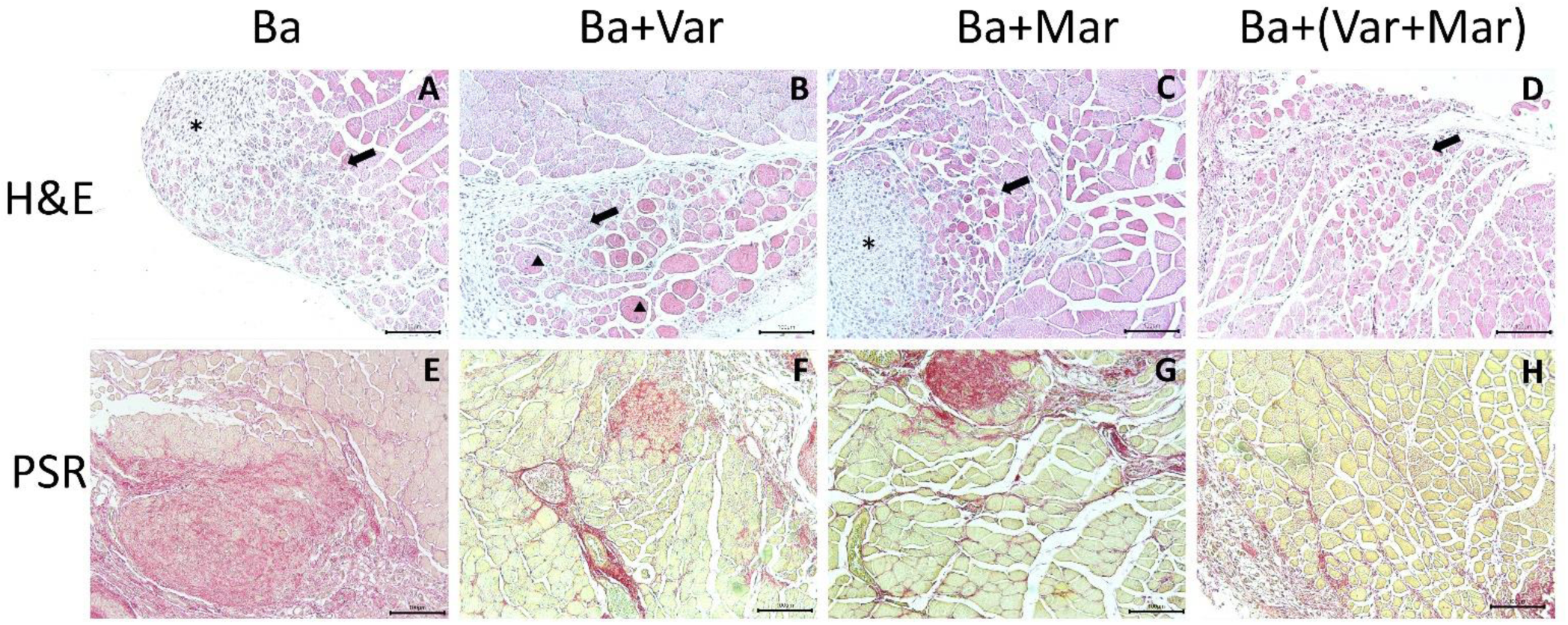
Light micrographs of sections of mouse gastrocnemius muscles 14 days after injection of venom followed by either PBS or inhibitors. CD-1 mice received an i.m. injection of 50 µg *B. asper* venom in 50 µL PBS. Then, 24 h after envenoming, different groups received an i.v. injection of either PBS (Ba, A and E), Varespladib (Ba+ Var, B and F), Marimastat (Ba + Mar, C and G) or Varespladib + Marimastat (Ba + Var + Mar, D and H). Mice were sacrificed 14 days after envenoming and tissue was processed and stained with hematoxylin-eosin (H & E) (upper images) or picrosirius red (PSR) (lower images). Deficient muscle regeneration is observed in sections from envenomed mice that did not receive the inhibitors, characterized by areas of regenerating fibers intermixed with areas of poor regeneration, with fibrosis and granulation tissue. Sections from tissues of envenomed mice treated with the inhibitors showed evidence of improved muscle regeneration and less extent of fibrosis. Arrows represent areas of regenerating muscle fibers with centrally-located nuclei, asterisks represent areas of poor regeneration and triangles represent unaffected muscle fibers. Bar represents 100 µm.

In order to provide a quantitative assessment of these observations, we evaluated the extent of regeneration by assessing the number of regenerating muscle fibers per muscle and the diameter of the regenerative fibers. All treatments with inhibitors induced a significant improvement in skeletal muscle regeneration 14 days after envenoming, as judged by the number of regenerating muscle cells per muscle, i.e., fibers with centrally-located nuclei; the highest effect was observed when the two inhibitors were combined (Fig 2A). On the other hand, no significant differences were observed in the diameter of regenerating fibers between treatments, except for the envenomed group treated with Marimastat which showed a significant increase in diameter of regenerating fibers as compared to samples from envenomed mice that did not receive inhibitors (Fig 2B). Regarding fibrosis, assessed by staining with picrosirius red, tissue from envenomed mice receiving Varespladib, Marimastat, or Marimastat + Varespladib showed a significant reduction in the extent of fibrotic areas of the muscle as compared to samples from mice receiving venom and no inhibitors (Fig 2C).

**Figure 2.**
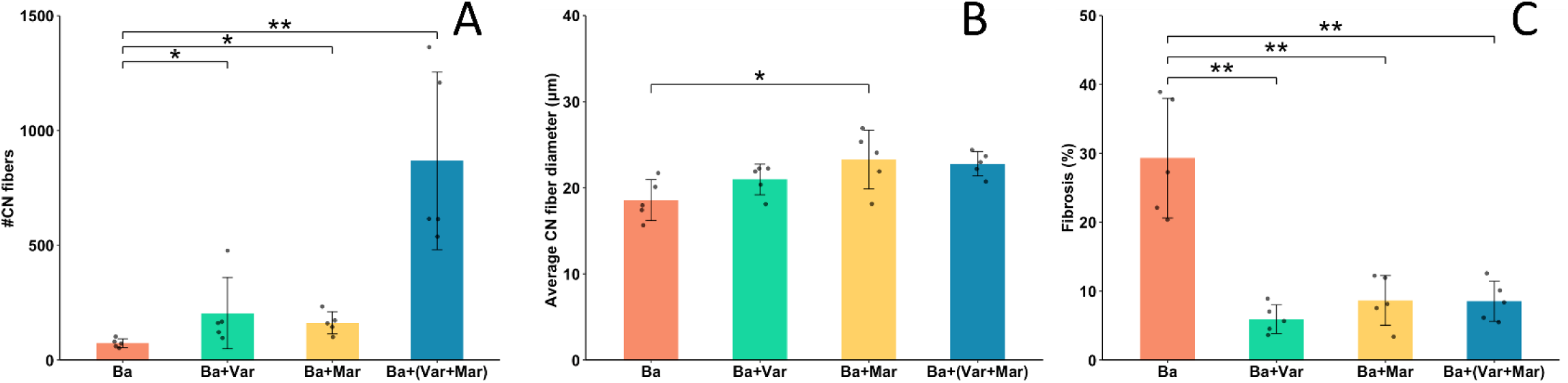
Quantitative histological assessment of skeletal muscle regeneration 14 days after injection of *B. asper* venom, followed 24 h later by the i.v. injection of either PBS (Ba), Varespladib (Ba + Var), Marimastat (Ba + Mar), or Varespladib + Marimastat (Ba + Var + Mar). Tissue was processed and sections were stained with hematoxylin-eosin or picrosirius red. (A) Number of fibers with centrally-located nuclei (CN) in the whole muscle section. (B) Diameter of all muscle fibers having centrally-located nuclei (CN). The diameter of normal muscle fibers in non-envenomed samples injected with PBS was 40.7 µm ± 8.3 (n = 400). (C) Percentage of the total area of muscle showing predominance of picrosirius red staining. Statistically significant differences were determined by negative binomial regression (A), one-way ANOVA with Tukey post-hoc test (B), and general linear model (GLM) with gamma distribution (C). Values represent mean ± S.D. (n = 5). *p < 0.05; **p < 0.001.

#### 2.2.2. Observations in samples collected 28 days post-envenoming

When the analyses were carried out in tissue samples collected 28 days after envenoming, a significant increase in the wet weight of the injected gastrocnemius muscle was observed in envenomed mice that received Marimastat, as compared to those that did not receive inhibitors, whereas no significant differences were observed with the other treatments. The weights of muscles from all envenomed groups were significantly different from that corresponding to mice receiving PBS and no venom, with the exception of the envenomed group treated with Marimastat (Supplementary figure S2).

Histological assessment of muscle tissue injected with venom and receiving PBS instead of inhibitors inhibitors showed a mixture of areas of regenerative muscle fibers of small diameter and areas of poor regeneration characterized by fibrotic tissue and regions showing apparent calcification. In the cases of samples from envenomed mice treated with the inhibitors, especially those receiving Marimastat or Marimastat + Varespladib, there was a better regenerative landscape, with centrally-nucleated muscle fibers intermixed with fewer areas of fibrosis. Assessment of fibrosis by picrosirius red staining evidenced an apparent reduction in interstitial collagen in all samples, as compared to samples collected at 14 days. However, tissue from envenomed mice not treated with the inhibitors showed a higher extent of picrosirius staining when compared to samples from envenomed mice that received the inhibitors, especially Marimastat or Marimastat + Varespladib (Fig. 3).

**Figure 3.**
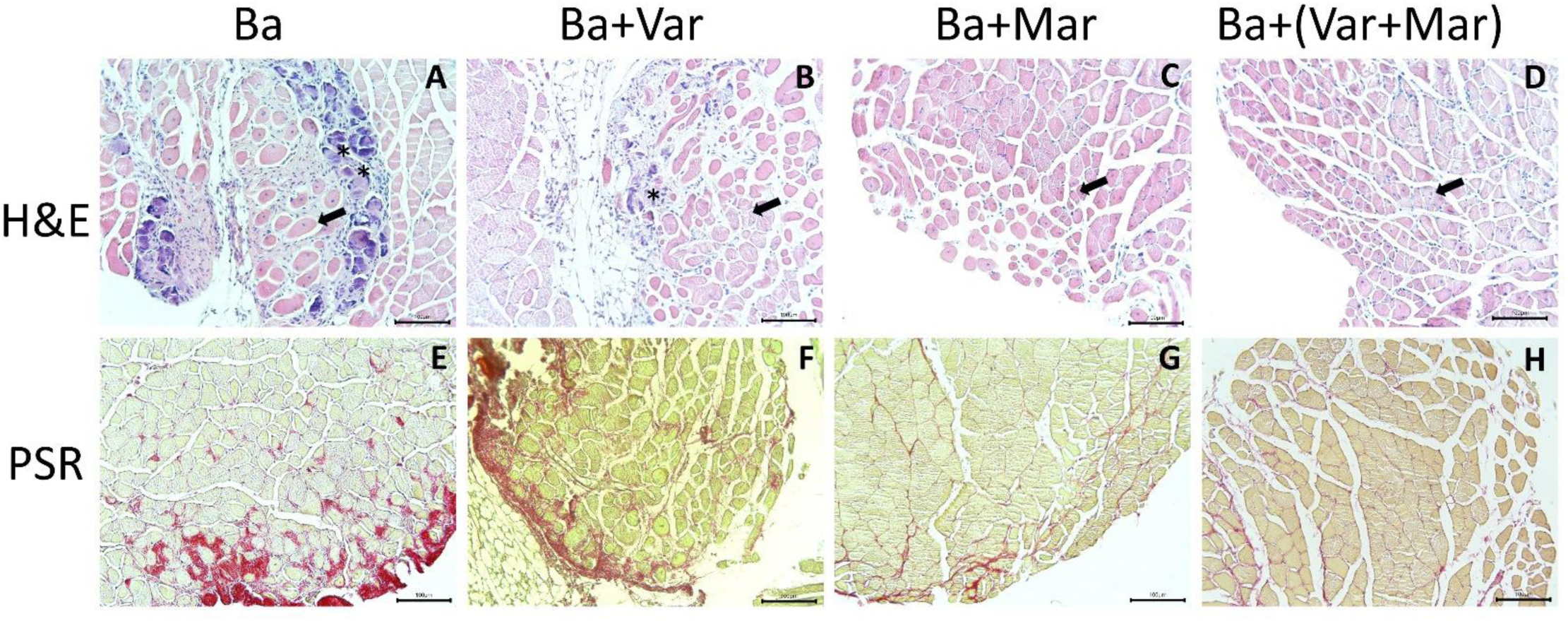
Light micrographs of sections of mouse gastrocnemius muscles 28 days after injection of venom followed by either PBS or inhibitors. CD-1 mice received an i.m. injection of 50 µg *B. asper* venom in 50 µL PBS. Then, 24 h after envenoming, different groups received an i.v. injection of either PBS (Ba, A and E), Varespladib (Ba + Var, B and F), Marimastat (Ba + Mar, C and G) or Varespladib + Marimastat (Ba + Var + Mar, D and H). Mice were sacrificed 28 days after envenoming and tissue was processed and stained with hematoxylin-eosin (H & E) (upper images) or picrosirius red (PSR) (lower images). Deficient muscle regeneration is observed in sections from envenomed mice that did not receive the inhibitors, characterized by areas of regenerating fibers intermixed with areas of poor regeneration and apparent calcification (basophilic staining). Sections from tissues of envenomed mice treated with the inhibitors, especially with Marimastat or Marimastat + Varespladib showed evidence of improved muscle regeneration, less extent of fibrosis and no calcification. Arrows represent areas of regenerating muscle fibers with centrally-located nuclei, whereas asterisks represent areas of poor regeneration. Bar represents 100 µm.

When a quantitative assessment of muscle regeneration was carried out in samples collected at 28 days, tissue from envenomed mice that received Marimastat or Marimastat + Varespladib showed a significantly higher number of regenerating fibers, as compared to mice injected with venom and no inhibitors. At this time interval, no significant improvement in regeneration was observed in the group of envenomed mice receiving Varespladib (Fig 4A). In agreement, a significant higher diameter of regenerating fibers was observed in samples of envenomed mice treated with Marimastat or Marimastat + Varespladib, but not in those receiving Varespladib only, when compared to samples from envenomed mice that did not receive inhibitors (Fig 4B). The extent of intramuscular fibrosis, as judged by picrosirius red staining, was reduced in all samples when compared to those collected at 14 days. When compared to samples from envenomed mice not receiving inhibitors, the percentage of picrosirius red-positive areas was reduced in samples from mice treated with Marimastat or Marimastat + Varespladib (Fig 4C).

**Figure 4.**
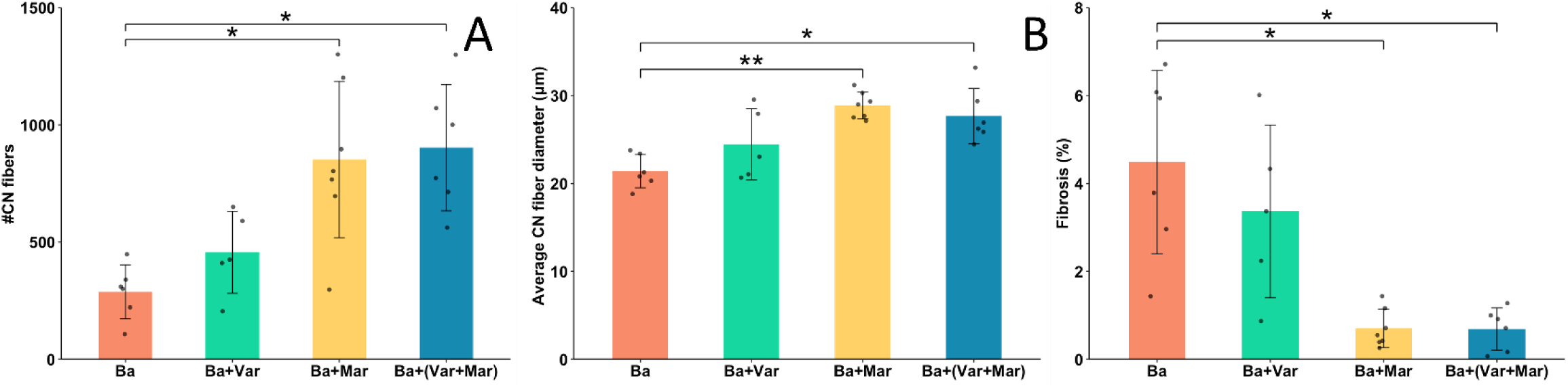
Quantitative histological assessment of skeletal muscle regeneration 14 days after injection of *B. asper* venom, followed 24 h later by the i.v. injection of either PBS (Ba), Varespladib (Ba + Var), Marimastat (Ba + Mar) or Varespladib + Marimastat (Ba +Var + Mar). Tissue was processed and sections were stained with hematoxylin-eosin or picrosirius red. (A) Number of fibers with centrally-located nuclei (CN) in the whole muscle section. (B) Diameter of all muscle fibers having centrally-located nuclei (CN). (C) Percentage of the total area of muscle showing predominance of picrosirius red staining. Statistically significant differences were determined by one-way ANOVA with Tukey post-hoc test (A, B) and general linear model (GLM) with gamma distribution (C). Values represent mean ± S.D. (n = 5-7). *p < 0.01; **p < 0.001.

## 3. Discussion

Our study explored whether the administration of Varespladib and Marimastat, or a combination of them, improves skeletal muscle regeneration after myonecrosis induced by *B. asper* venom. These inhibitors were administered by the i.v. route 24 h after venom injection, a time when acute muscle damage has occurred and the inflammatory and regenerative processes are under way [15–17]. Thus, the potential beneficial effects of these inhibitors are not due to inhibition of venom-induced acute tissue damage, which has already occurred by 24 h, but instead to the inhibition of venom or endogenous enzymes present in the tissue during the early stages of muscle regeneration. It has been shown that traces of venom remain in the necrotic muscle even days after envenoming [10], having deleterious effects on muscle regeneration, as evidenced by the inhibitory action of muscle homogenates, obtained several days after venom injection, on myoblast replication and fusion *in vitro*, and the abrogation of this inhibitory effect when homogenates are incubated with antivenom [10,11].

Our approach allowed a qualitative and quantitative assessment of muscle regeneration by using various parameters in tissue samples collected at 14 and 28 days. These times were selected on the basis of previous studies showing that, at these intervals, regeneration is well advanced in models of myonecrosis induced by myotoxic PLA_2_s in which there is a good regenerative response, while it is partially deficient after myonecrosis induced by complete *B. asper* venom [5–7].

Varespladib and Marimastat were selected as inhibitors because they inhibit snake venom PLA_2_s, in the case of Varespladib, and SVMPs, in the case of Marimastat, in a variety of snake venoms [18–23]. Varespladib reduced the extent of muscle damage and improved muscle regeneration after injection of the venom of *Deinagkistrodon acutus* [22]. Moreover, Varespladib and Prinomastat, another metalloproteinase inhibitor, abrogate atrophy induced by the venom of *D. russelii* on myotubes in culture [12]. Thus, since homogenates from muscle injected with a myotoxic PLA_2_ and a hemorrhagic SVMP from *B. asper* venom were shown to inhibit myoblast fusion in culture [10], it was hypothesized that inhibiting these toxins in the regenerating muscle would improve the overall regenerative process.

Marimastat, and especially the combination of both inhibitors, showed the strongest pro-regenerative effect. It was of interest that the combination induced a marked increase in the number of regenerating fibers by 14 days, whereas by 28 days the treatments based on Marimastat or the combination of inhibitors resulted in a similar improvement. These findings suggest that SVMPs remaining in the tissue exert a stronger inhibitory role of the regenerative process than PLA_2_s. However, the fact that the combination of Marimastat and Varespladib induced a higher extent in regeneration at 14 days, as judged by the number of regenerating cells in the damaged muscle, suggests that they may be acting synergistically in promoting the early stages of regeneration. On the other hand, the observation that the number of regenerating fibers is similar at 14 days in samples of mice treated with the combination of inhibitors and at 28 days in samples of mice treated with Marimastat or with the combination, suggests that muscle regeneration reaches a plateau and that it is accelerated when the two inhibitors are used together.

Varespladib was originally designed to inhibit endogenous PLA_2_s for the treatment of rheumatoid arthritis, sepsis and acute coronary syndrome [24–26] and Marimastat for inhibiting endogenous matrix metalloproteinases (MMPs) in vascular disease and malignancies [27, 28]. Thus, the possible inhibition of endogenous enzymes in muscle tissue having a role in muscle repair and regeneration cannot be discarded. MMPs are known to play key roles in muscle repair and regeneration [29, 30, 31] and their activities increase in muscle injected with PLA_2_ and SVMP from *B. asper* venom [32]. MMPs also play a role in the regulation of muscle resident fibro/adipogenic precursors which proliferate and give rise to adipocytes in various muscle pathologies [33]. Thus, the inhibitors used in our study may be working not only against snake venom enzymes but also against endogenous enzymes, an issue that requires further studies.

The mechanisms by which *B. asper* venom PLA_2_s and SVMPs affect muscle regeneration remain unknown. Myotoxic PLA_2_s are cytotoxic to myoblasts and myotubes in culture [34, 35], and apoptotic muscle regenerative cells have been described in regenerating muscle after myonecrosis induced by *B. asper* venom [7]. On the other hand, SVMPs affect the myogenic satellite cells [36] and degrade the basement membrane that surrounds muscle fibers, which plays a key role in the regenerative response [36,37]. Moreover, since revascularization is relevant during muscle regeneration, the damaging effect of hemorrhagic SVMPs on the microvasculature [7, 38] may also affect angiogenesis. In our experimental setting we administered the inhibitors by the intravenous route 24 h after venom injection, a time when activation and proliferation of myogenic cells in damaged muscle is underway. Thus, the inhibitors are likely to block venom SVMPs and PLA_2_s by the time key steps in the regenerative process are taking place. It would be relevant to test other protocols such as using repetitive administrations and variable doses of the inhibitors during the first week after the onset of myonecrosis, when additional stages in the muscle regenerative process are taking place, in order to assess whether regeneration can be improved with inhibitors even further.

In conclusion, our findings provide clues to understand one of the mechanisms responsible for the poor muscle regeneration after myonecrosis induced by viperid snake venoms, i.e., the deleterious effects of venom toxins remaining in the tissue during the first stages of muscle regeneration which may affect the regenerative process in various ways. The beneficial role of Marimastat, or the combination of Marimastat and Varespladib, suggests that SVMPs may be involved in such effect by directly damaging myogenic precursors or hydrolyzing extracellular matrix components necessary for regeneration. In addition, our results also suggest that the administration of inhibitors of metalloproteinases, even after muscle damage has occurred, may represent a promising therapeutic avenue to improve muscle regeneration after myonecrosis induced by viperid venoms. It would be relevant to assess whether such enhanced regenerative response also occurs after myonecrosis induced by other snake venoms.

## 4. Materials and methods

### 4.1. Venom and inhibitors

The venom of the snake *Bothrops asper* was used throughout this study. Venom was a pool obtained from more than 40 adult specimens collected in the Pacific region of Costa Rica and maintained at the serpentarium of Instituto Clodomiro Picado. After lyophilization, venom was stored at -20 °C. Venom solutions were prepared in 0.12 M NaCl, 0.04 M phosphates, pH 7.2 (PBS) immediately before use. The inhibitor Varespladib (LY315920) was provided by Ophirex, Inc. (Corte Madera, CA, USA) via ChemiTek (Indianapolis, IN, USA), and Marimastat (BB-2516) was obtained from Sigma-Aldrich (St. Louis, MO, USA).

### 4.2. Experimental animals

Throughout this study, mice (CD-1 strain, 18-20 g body weight of both sexes) were used. Mice were provided by the Bioterium of Instituto Clodomiro Picado. They were housed in plastic cages and kept under controlled temperature, and a 12:12 h light-dark cycle, and were provided with water and food *ad libitum*. The protocol of experiments involving animals was approved by the Institutional Committee for the Care and Use of Animals (CICUA) of the University of Costa Rica (approval code CICUA-17-2023).

### 4.3. Quantification of muscle tissue necrosis induced by B. asper venom

For the assessment of the extent of muscle tissue necrosis induced by *B. asper* venom at 24 h, a group of eight mice received an intramuscular (i.m.) injection, in the right gastrocnemius, of 50 µg venom dissolved in 50 µL PBS. A control group of mice received 50 µL PBS under otherwise identical conditions. Thirty minutes before venom injection, the analgesic tramadol was administered by the subcutaneous route at a dose of 50 mg/kg to reduce pain associated with tissue damage. At 24 h post-injection, mice were sacrificed by cervical dislocation. Immediately afterwards, the right gastrocnemius muscles were dissected and submerged in Zinc Formalin Fixative (Sigma-Aldrich, St Louis, MO, USA) for 24 h at 4°C. The tissue was dehydrated with serial ethanol and xylol and embedded in paraffin, and tissue sections of 4 µm thickness were obtained from the mid-belly section of the muscle.

Tissue was oriented to obtain transversal sections, deparaffinized with xylene, hydrated in distilled water and stained with hematoxylin and eosin for microscopic evaluation. The totality of each cross section of muscle tissue was captured in several images using a light microscope Delphi-X Observer (Euromex, Arnhem, The Netherlands) with a camera. The total muscle area and the areas of necrotic muscle, i.e., areas with fibers showing hypercontraction of myofibrillar material, were measured, and the extent of muscle damage was expressed as the percentage of the areas of myonecrosis relative to the total muscle area.

### 4.4. Effect of Varespladib and Marimastat on muscle regeneration after myonecrosis induced by B. asper venom

For the evaluation of the effect of Varespladib and Marimastat on muscle regeneration, groups of 5-7 mice received an i.m. injection of 50 µg venom, dissolved in 50 µL PBS, as described above. Twenty-four hours after venom injection, mice received an intravenous (i.v.) administration of 100 µL of either PBS, Varespladib (0.8 mg/mL), Marimastat (0.8 mg/mL), or a combination of both inhibitors injected separately (each one at 0.8 mg/mL). Inhibitors were dissolved first in DMSO (20 mg/mL) and diluted in PBS to reach the final concentration used. Mice were sacrificed by cervical dislocation 14 and 28 days after venom injection, and the right gastrocnemius muscles were dissected for tissue processing, as described above. In samples collected 28 days after venom injection, the wet weights of the injected gastrocnemius muscles were determined before tissue processing.

For the analysis of muscle regeneration, samples of tissue collected 14 and 28 days after envenoming were analyzed. The extent of muscle regeneration was expressed in two ways: (a) the total number of regenerating muscle fibers in the total area of the muscle section, and (b) the diameters of regenerating muscle fibers. Regenerating muscle fibers were identified as those having a centrally-located nuclei, which is a morphological feature of regenerating fibers that is maintained for a prolonged period of time after the onset of necrosis induced by a snake venom toxin [39]. For these quantitative assessments, the Delphi-X Observer (Euromex) with the Image Focus Alpha (Euromex) software was used.

For detection of fibrotic areas in muscle, 4 µm muscle sections of tissue processed as described above were stained for one hour with a 0.1% (wt/vol) Sirius red, C.I. 35780 (Polysciences Inc., Warrington, PA, USA) dissolved in saturated aqueous picric acid (Sigma-Aldrich), following the protocol of Dapson et al. [40]. The slides were dehydrated with three changes of 100% of ethanol, cleared in three changes of xylene 100% and mounted in Dako Mounting Medium (Statlab Medical Products, McKinney, TX, USA). Micrographs from muscles were captured using a Delphi-X Observer (Euromex). The total area of the muscle section was measured, as well as the areas in which fibrosis predominated, i.e., areas inside muscle fascicles stained positively with picrosirius red. Staining corresponding to dense fibrous connective tissue located in regions between muscle fascicles was not considered. The extent of fibrosis was expressed as the percentage of the area with predominant staining with picrosirius red relative to the total area of the tissue examined. The collected images were analyzed using the software Image Focus Alpha (Euromex) and ImageJ (NIH, Bethesda, MD, USA). In the study of hematoxylin-eosin and picrosirius stained sections, samples were anonymized and the analyses were carried out by two blinded observers.

### 4.5. Statistical analysis

Graphics for the figures were made using the program RStudio version 4.5.1 (2025-06-13) (R foundation). Differences between experimental groups were assessed using a one-way ANOVA followed by Tukey *post-hoc* test for normally distributed data. For non-normally distributed data, comparisons were performed using generalized linear model (GLM) with gamma or negative binomial distributions, as appropriate; p values <0.05 were considered statistically significant.

## Acknowledgements

The authors thank Alejandro Navarro for his collaboration in the preparation of samples. This study was supported with funding by the National Institutes of Health Fogarty International Center grant number D43TW011403 entitled ‘International Training Program in Environmental Health over the Lifespan’ (Claudio L & van Wendel de Joode B, PIs), a grant assigned to the Icahn School of Medicine at Mount Sinai and the Universidad Nacional, Costa Rica, .and Vicerrectoría de Investigación, Universidad de Costa Rica (project C4600).

## Supplementary files

**Supplementary figure S1.**
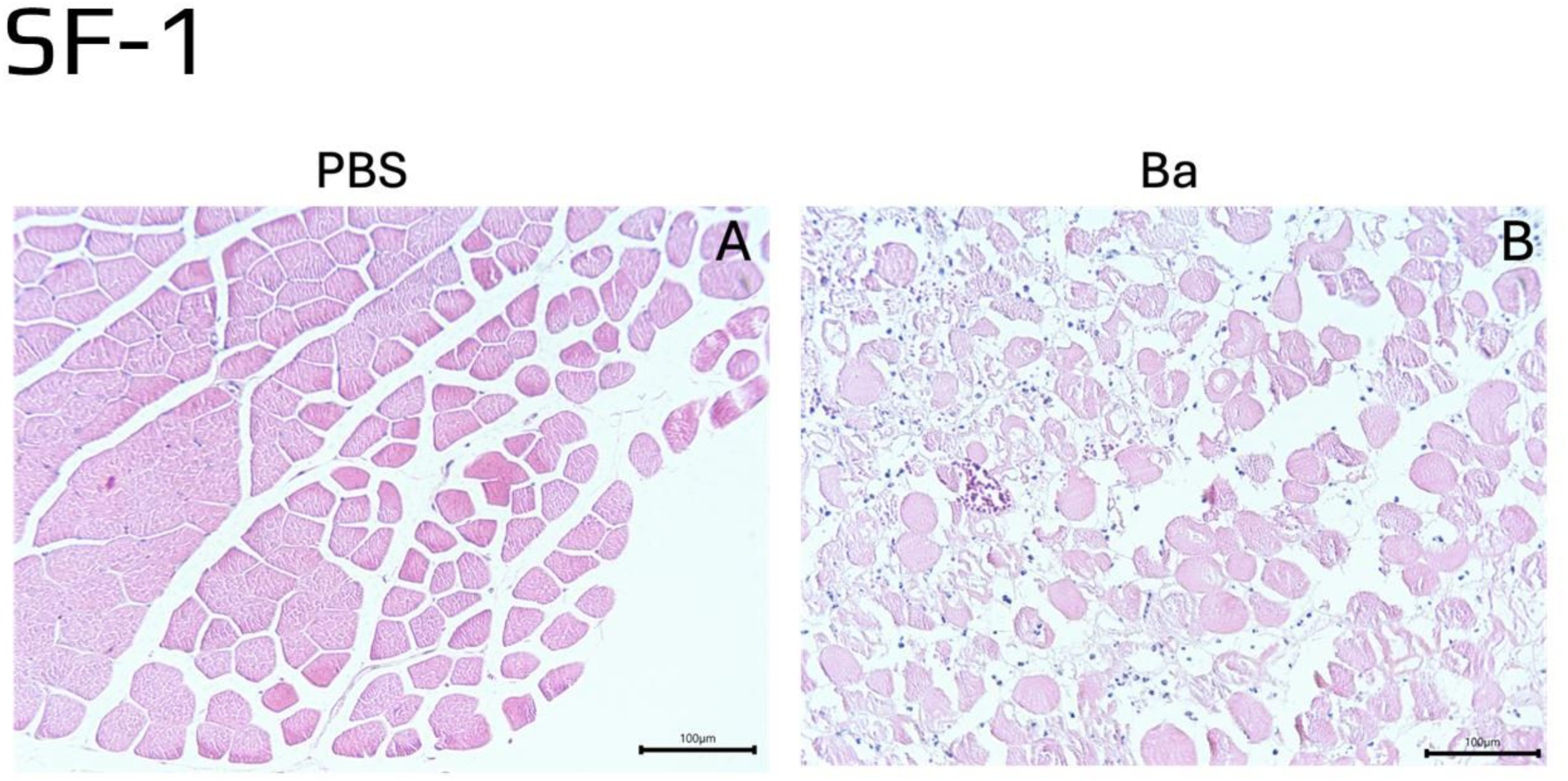
Light micrographs of sections of gastrocnemius muscles of mice receiving an i.m. injection of either PBS (A) of 50 µg *B. asper* venom (Ba), dissolved in 50 µL PBS. Tissue samples were obtained 24 h after injection and processed for embedding in paraffin and staining with hematoxylin-eosin. Notice the normal histological pattern in A and the prominent necrosis of muscle fibers in B. Bars represent 100 µm.

**Supplementary figure S2.**
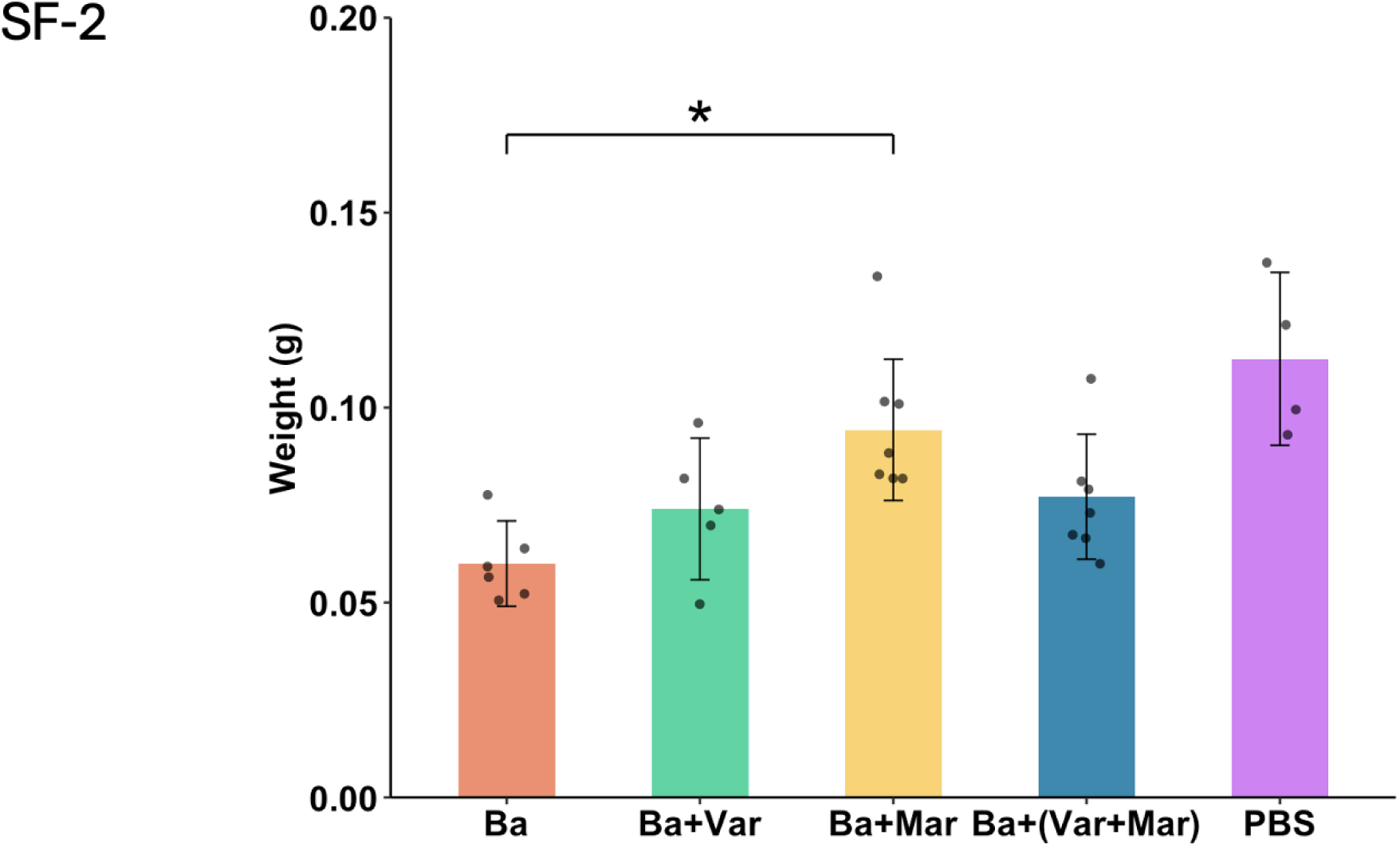
Wet weight (g) of the right gastrocnemius muscle of CD1 mice, at 28 days after intramuscular injection of 50 µg *B. asper* venom, dissolved in 50 µL PBS, followed, 24 h later, by the intravenous administration of either PBS (Ba), Varespladib (Ba + Var), Marimastat (Ba + Mar) or Varespladib + Marimastat (Ba + Var + Mar). The bar of PBS corresponds to non-envenomed mice receiving PBS. Wet weight of envenomed mice receiving Marimastat was significantly higher than that of envenomed mice not receiving inhibitors (*p < 0.05). Statistical analysis was performed using one-way ANOVA with Tukey *post-hoc* test. The weight of the muscle of mice injected with PBS only was significantly different from: Ba (p <0.001), Ba+Var (p <0.05) and Ba+(Var+Mar) (p <0.05). No significant difference was observed between the treatments Ba+Mar and PBS (p = 0.44).

